# Microbiomes from feces *vs.* gut in aquatic vertebrates: distinct community compositions between substrates and preservation methods

**DOI:** 10.1101/651612

**Authors:** Sten Anslan, Huan Li, Sven Künzel, Miguel Vences

**Affiliations:** Zoological Institute, Technische Universität Braunschweig, Mendelssohnstr. 4, 38106 Braunschweig, Germany; Institute of Occupational Health and Environmental Health, School of Public Health, Lanzhou University, Lanzhou 730000, China; Department of Evolutionary Genetics, Max Planck Institute for Evolutionary Biology, Plön, Germany

**Keywords:** *Nanorana*, *Gymnocypris*, *Triplophysa*, gut microbiota, fecal samples, metabarcoding, 16S rRNA

## Abstract

Sample type and preservation methods are likely to influence microbiome analysis results. Relatively few studies have explored the differences between feces and gut as well as ethanol-stored and frozen samples. Here, we sampled the same individuals of three aquatic vertebrates from the Qinghai-Tibetan Plateau non-invasively for feces, and subsequently for hindgut through dissection. Our study species, two fishes (*Gymnocypris* cf. *namensis* and *Triplophysa* sp.) and one amphibian (tadpoles of *Nanorana parkeri*), were all collected at the same time and site. Gut and fecal samples were stored in ethanol, and additionally, part of the gut samples were frozen, but temporarily thawed during transport as it often happens under difficult field conditions. Our results showed that both substrate (gut content *vs*. feces) as well as preservation method can influence the analysis of intestinal microbiomes. Frozen gut samples strongly differed from ethanol-stored samples, and especially in *Nanorana* most frozen samples were dominated (in relative abundance) by a set of Proteobacteria OTUs that were completely absent from the ethanol-stored samples. This blooming of contaminant bacteria occurred after less than 12 h of thawing, thus caution should be taken when constancy of cold temperatures cannot be maintained in the field for sample preservation purposes. Among ethanol-stored samples, bacterial communities from feces differed from those recovered from guts, but in part recovered similar patterns, such as a higher bacterial richness in the more herbivorous *Nanorana* tadpoles. Although our results argue against combining gut and fecal samples in analyses of host-specific microbiome differences, they also confirm that non-invasive sampling of feces can provide useful information of gut microbiomes in aquatic vertebrates, which may be important especially when working with endangered species.

## Introduction

Many studies have targeted the composition, dynamics and highlighted the health relevance of the human gut microbiome [e.g. 1, 2]. Likewise, animal-associated microbial communities play important roles in the biology and health of their hosts [3], including food degradation and energy harvest [4], immunity regulation [5], and physical development [6]. These studies are typically based on high-throughput sequencing (HTS) of short 16S rRNA gene amplicons, where significantly different composition and diversity patterns of the host-associated microbiomes are driven by host taxonomy, ecology and environment [7-12].

While the intestinal microbiome of mammals has been the subject of numerous studies [13-16], research on microbiomes associated with aquatic vertebrates is still in its infancy. Fish microbiomes have been studied in the context of aquaculture, but work on wild fish is relatively rare [17-19]. Research on the gut microbiome of amphibians is even less frequent. Work on this vertebrate group has typically focused on the cutaneous microbiome [20, 21], often in the context of its effects on pathogenic fungi causing amphibian declines [22, 23]. Work on the amphibian gut microbiome is scarce [24-29] but of substantial biological interest especially in frogs, given the drastic restructuring of the gut in the metamorphosis from a largely herbivorous tadpole to an exclusively carnivorous frog [10, 30].

Understanding which factors influence microbiomes at local and global scale is of importance to unravel general biogeographic and macro-ecological trends [21, 31, 32] and host-microbiome interactions [12], including diseases. Sampling the microbiome of the intestine is a moderately to strongly invasive approach, especially in small animals where it requires killing the studied individuals. Instead, samples of fresh feces are often used and the fecal microbiome is considered as representative of the gut microbiome [e.g. 12, 33]. However, indications exist that fecal characteristics influence the composition and richness of detected microbiota [34]. Therefore, it is uncertain to which extent the fecal microbiome may serve as proxy for the gut microbiome across hosts, and whether communities from feces and gut samples are fully comparable [e.g. 35]. In particular, only few studies address this question in tadpoles [36] and other aquatic animals, although several studies used non-invasive collection of fecal samples from fishes to study the composition of intestinal microbiomes [e.g. 37].

For large-scale analyses, especially meta-analyses of data sets originating from a diversity of sources, comparability of data is a basic prerequisite. This refers not only to laboratory methods but also extends to sampling and sample preservation [38]. Thus, a further factor influencing the inference of microbiome composition from high-throughput sequencing of amplicons is the method of sample preservation. Freezing samples at −20°C immediately upon collection has been defined as the gold standard to ensure the microbial community does not change until DNA extraction [39]. Keeping samples uninterruptedly at this temperature is however not always possible under difficult field conditions, which might lead to alterations of the microbial community composition during episodes of thawing [40].

In our study, we tested the effect of sampling substrate, feces *vs*. hindgut, to characterize the gut microbiota of tadpoles of one species of frog (*Nanorana parkeri*), and of two species of fish (*Gymnocypris* cf. *namensis* and *Triplophysa* sp.). In addition, we compared the similarity of the detected microbiota using freezing *vs*. ethanol (EtOH) preservation method of the samples from the same specimens. The sample types per specimen included EtOH-stored feces, EtOH-stored gut and frozen gut samples. As feces samples are widely considered to reflect gut microbiota, we expected similar richness and community patterns from the EtOH-stored feces and gut samples. In this study, the frozen gut samples were exposed to thawing episodes during sample transport from the remote location, thus we predicted a skewed microbial community composition in comparison with EtOH-stored gut samples and tested the extent and constancy of this effect.

## Methods

### Study site and sampling methods

Specimens of fish and tadpoles were collected on 2^nd^ of July 2018 from the central Qinghai-Tibetan Plateau, China, in a small tributary stream and a small pond directly nearby the stream, in the area of Lake Nam Co (N30.82840°, E91.06397°). Eight specimens of *Gymnocypris* cf. *namensis, Triplophysa* sp. and twelve tadpole specimens of *Nanorana parkeri* were collected using dip nets and placed individually into sterile Whirl-Pak R bags together with a small amount of clean water from the respective water bodies. A pond water control sample was collected by dipping a sterile swab into the water, placing it into a cryotube and freezing it. Tadpoles were in Gosner stages 26-30 whereas the fishes were likely sub-adults, at total lengths of approximately 50-100 mm. Specimens were kept overnight (ca. 14 h) in the bags, anesthetized with tricaine methanesulfonate (MS222; Sigma-Aldrich, St. Louis, MO, USA) solution and subsequently overdosed using MS222 (a procedure not requiring approval by an ethics committee at TU Braunschweig). We collected the feces accumulated in the bags using pipettes, as well as a portion of hindgut of every individual. In summary, we obtained three replicate samples per specimen: (1) feces, stored in 100% EtOH; hindgut, stored in 96% EtOH; (3) hindgut, frozen at −20° C right after collection. Samples were frozen upon collection, but as typical for suboptimal fieldwork conditions, underwent two thawing-freezing cycles during transport to the lab, with temperatures of 10-15°C for limited times <12 h. In the laboratory, all samples were stored at −20° C until further processing.

### Molecular analysis

Following the manufacturer’s instructions, DNA was extracted using DNeasy PowerSoil Kit (QIAGEN, Germany). PCR was performed using the forward primer 515F (5’-GTGCCAGCMGCCGCGGTAA-3’) and reverse primer 806R (5’-GGACTACHVGGGTWTCTAAT-3’) to target the V4 region of 16S rRNA gene [41]. The used primer and tag combinations for each sample are specified in Supplementary data 1. For PCR, the 25 µl mixture per sample comprised of 2 µl DNA (3 µl for repeated samples), 0.5 µl each of the primer, 4 µl 5× HOT FirePol^®^ Blend Master Mix (Solis BioDyne, Tartu, Estonia) and 17 µl sterile dH_2_O. PCR was carried out in two replications in the following thermocycling conditions: an initial 15 min at 95 ° C, followed by 35 cycles of 94 ° C for 45 s, 50 ° C for 60 s, 72° C for 90 s, and a final cycle of 10 min at 72° C. PCR products per sample were pooled and their relative quantity was estimated during gel electrophoresis of 5 µl DNA sample on 1% agarose gel (15 min). Based on the gel band strength, all PCR products were pooled at approximately equimolar concentration. The DNA library was purified using Favor-Prep™ Gel/PCR Purification Kit (Favorgen-Biotech Corp., Austria), following the manufacturer’s instructions. Sequencing was performed on the Illumina MiSeq (2×250) using MiSeq Reagent Kit v2. Steps of DNA extraction, PCR and sequencing included both negative and positive controls (Supplementary data 1). Sequencing data sets have been deposited in the Sequence Read Archive (SRA): BioProject ID PRJNA533915.

### Bioinformatics

The paired-end sequence data was processed in QIIME [v1.9.0; 42] using the Environmental Microbiome and Bioinformatic Analysis Platform of School of Public Health in Lanzhou University. Data analysis methods were described previously [16]. Briefly, paired-end sequences were joined using Flash software [v1.2.8; 43]. Those sequences with length < 300bp, average base quality score < 30 or ambiguous bases, were removed for the downstream analysis. Uchime algorithm [44] was used to filter out potential chimeric reads. The filtered sequences were clustered into operational taxonomic units (OTUs) at a 97% identity threshold using UCLUST algorithm [45]. Taxonomy was assigned using the Ribosomal Database Project classifier [46]. OTUs not classifying to bacteria (Eukaryote and Archaea lineages) were removed. To account for the unequal sequencing depth, each sample was rarefied to the same number of reads (5654 sequences). The latter led to discarding one of the frozen gut sample of *Nanorana* from the downstream analysis. OTU table was further filtered to remove singleton OTUs and low-abundance read records of OTUs per sample (< 10 reads). After these step, total of 150 (out of total 988 OTUs) Cyanobacterial OTUs were removed from the analysis as these taxa likely do not represent true gut microbiota (e.g. Nostocophycideae, Synechococcophycideae, Oscillatoriophycideae [47]). The filtered OTU table used for the analyses is specified in Supplementary data 2.

### Statistical analysis

The effect of sample type (EtOH-stored feces, EtOH-stored gut and frozen gut) and species (*Gymnocypris, Triplophysa, Nanorana*) on log-transformed OTU richness were tested using factorial ANOVA (Type III SS) followed by Tukey HSD tests. The effect of these factors on the bacterial OTU composition was analyzed using PERMANOVA+ [48] with 9999 permutations (Type III SS) in PRIMER v6 [49]. For the PERMANOVA analysis and non-metric multidimensional scaling (NMDS) graphs, we used Hellinger-transformed Bray-Curtis as well as UniFrac distance (unweighted) OTU matrices. UniFrac distance and phylogenetic diversity (PD) were calculated by applying PhyloMeasures package [v2.1; 50] in R [51] using Maximum-likelihood based phylogenetic 16Sr RNA gene tree generated with RAxML [52]. We used indicator species analysis [v1.7.6; 53] to determine which OTUs are significantly (using 9999 permutations) associated with particular sample types.

## Results

Our analysis comprised samples of eight individuals for each of the two fish species, and 12 individual tadpoles. Replicate samples of EtOH-stored feces, EtOH-stored gut and frozen gut per specimen demonstrated high OTU richness variability, with an average of 2.13, 2.06 and 4.03-fold difference for *Gymnocypris, Triplophysa* and *Nanorana*, respectively (Fig. 1). Consequently, there was no strong positive correlation of the detected OTU richness between different sample types (except between *Triplophysa* EtOH-stored and frozen gut samples, and somewhat between *Nanorana* EtOH-stored feces and EtOH-stored gut samples; Supplementary data 3). Comparing the encountered pattern across the three species, OTU richness in the EtOH-stored samples was consistently highest in the *Nanorana* tadpoles as can be expected given their predominantly herbivorous diet, followed by *Gymnocypris*, and *Triplophysa* having lowest richness values (Fig. 2A). This pattern, however, was conspicuously reversed for the frozen samples, in which the *Nanorana* tadpoles had by far the lowest richness values, whereas no clear differences between the two fish species were appreciable (Fig. 2A). The phylogenetic diversity (PD) of the sample types exhibited comparable mean values, except frozen gut samples of *Triplophysa*, which demonstrated significantly lower PD (Fig. 2B).

**Figure 1.**
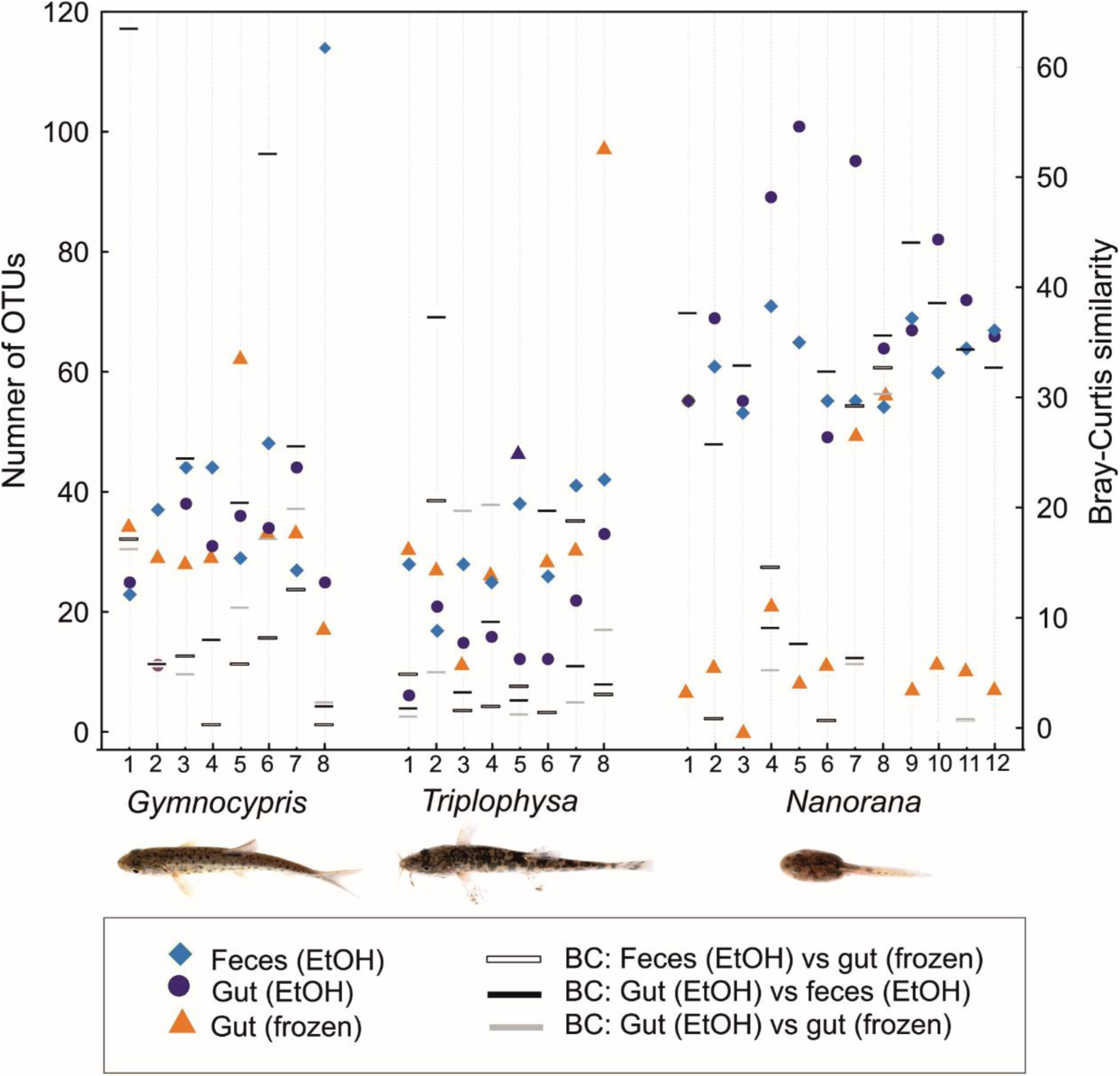
OTU richness (left y-axis) and Bray-Curtis (BC) similarity index (right y-axis) of each replicate sample per specimen. Missing BC index denotes 0 similarity between samples. Note that the frozen gut sample of *Nanorana* specimen 3 was discarded during rarefaction process, thus here denoted as 0 recovered OTUs.

**Figure 2.**
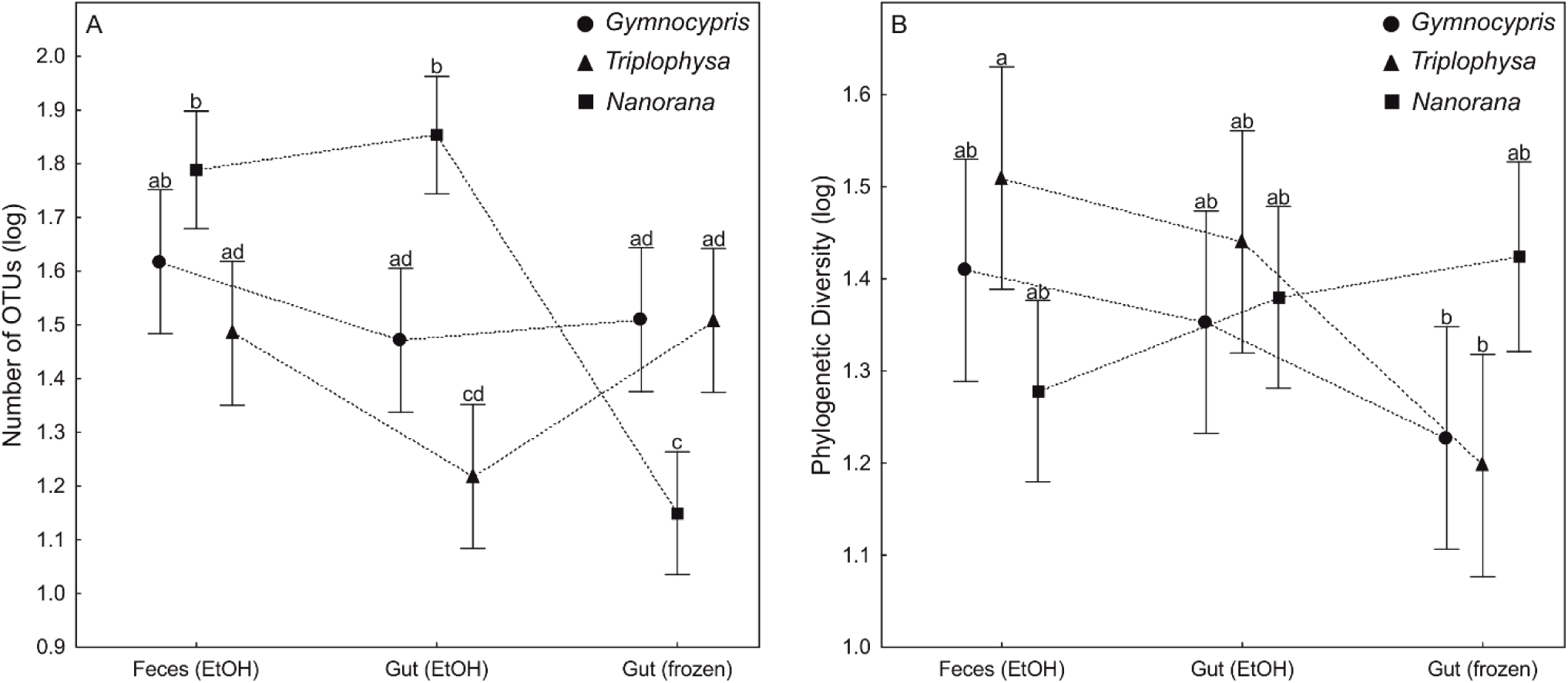
(A) OTU richness (log transformed) and B) phylogenetic diversity (log transformed) for EtOH-stored feces, EtOH-stored gut and frozen gut samples of *Gymnocypris* cf. *namensis, Triplophysa* sp., *Nanorana parkeri*. Whiskers denote 95% confidence intervals. Different letter combinations on top of whiskers denote significant difference between groups (α < 0.05) as based on Tukey HSD test.

Because of the species effect on bacterial community composition (pseudo-F_2, 82_ = 5.117, R^2^_adj_ = 0.102, P < 0.001), the following analyses were conducted separately for *Gymnocypris, Triplophysa* and *Nanorana*. In each case, the communities of detected microbiota were significantly different between sample types (EtOH-stored feces, EtOH-stored gut and frozen gut; Table 1). However, in *Gymnocypris* samples, the pairwise comparison revealed that this was not the case between EtOH-stored feces and EtOH-stored gut samples (Bray-Curtis matrix: P = 0.111; Table 1; Fig. 3e). Nonetheless, UniFrac distance matrix indicated pronounced difference of the latter (P = 0.007; Table 1; Fig. 3a). Although, the NMDS plots of *Triplophysa* samples (Fig. 3b, f) indicate differences in sample location (as well as in dispersion), the pairwise comparison showed no significant difference between EtOH-stored gut and frozen gut samples (Table 1). The most pronounced differences between sample types were observed in *Nanorana* samples (Table 1; Fig. 3c, g).

**Table 1.**
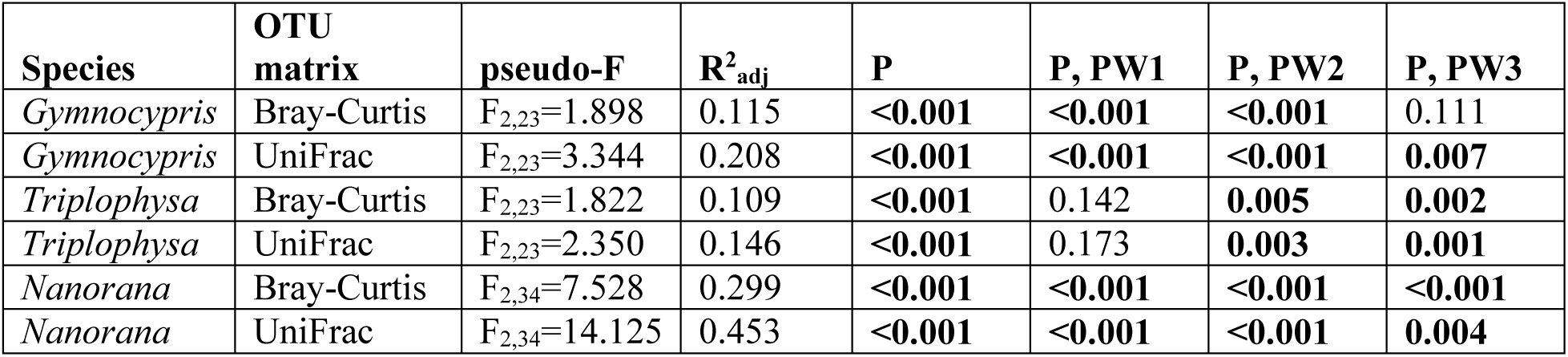
PERMANOVA results for *Gymnocypris, Triplophysa* and *Nanorana* samples with factor sample type (EtOH-stored feces, EtOH-stored gut and frozen gut). PW denote pairwise comparison between EtOH-stored gut and frozen gut (PW1), EtOH-stored feces and frozen gut (PW2), EtOH-stored feces and EtOH-stored gut (PW3) samples.

**Figure 3.**
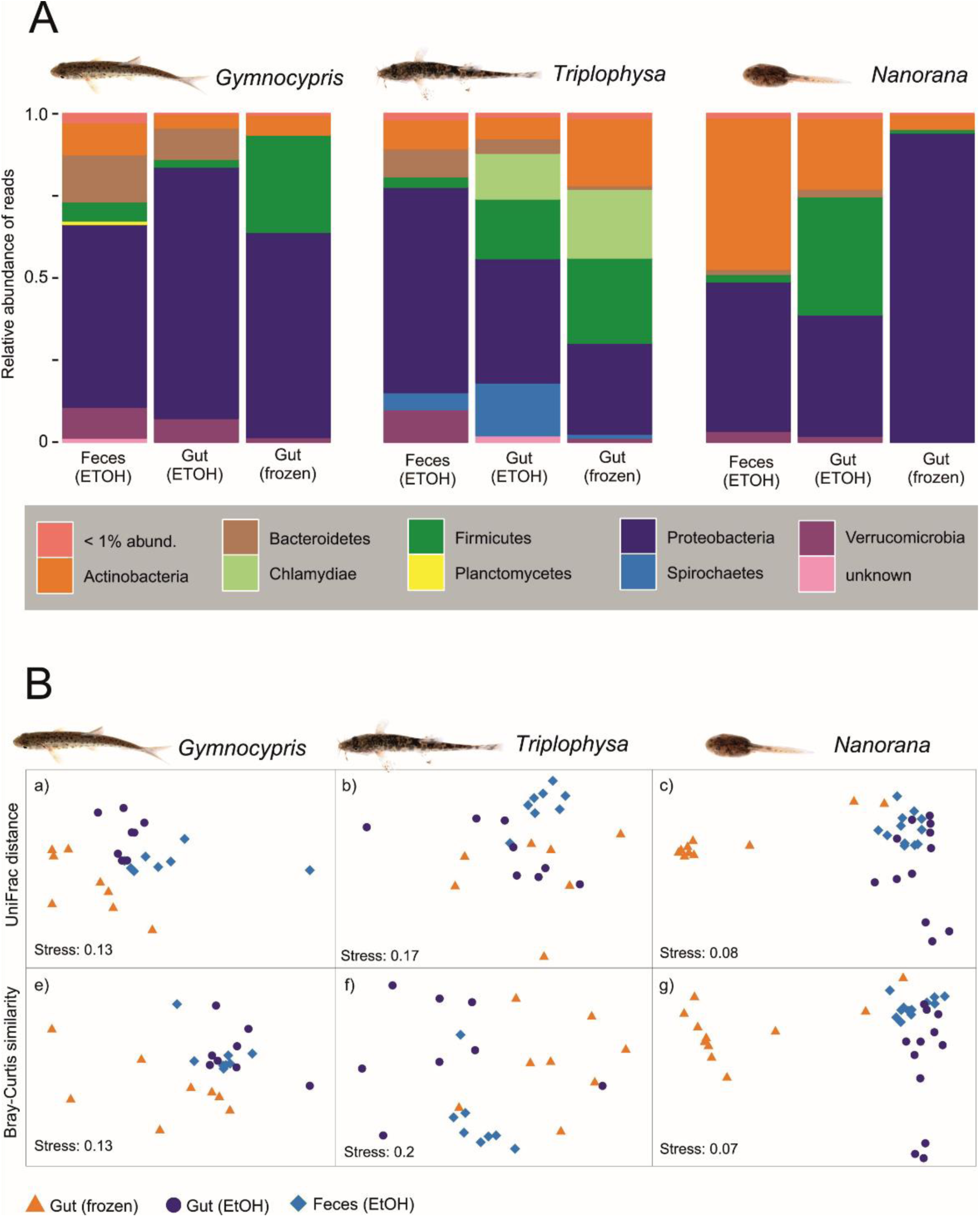
(A) Barplot showing relative abundance of reads assigned to bacterial phyla, summarized for each host species and treatment. (B) NMDS graphs as based on UniFrac distance (a-c) and Bray-Curtis similarity of bacterial community composition in *Gymnocypris* (a, e), *Triplophysa* (b, f) and *Nanorana* (c, g) samples.

Indicator species analysis revealed several OTUs specific to the sample type of each species (Supplementary data 4). Compared with frozen gut samples, consistenly more OTUs were assigned to be specific to EtOH-stored feces and gut samples. Moreover, the combination of EtOH-stored samples (feces + gut) showed to harbor many indicator OTUs, whereas no OTUs were assigned to be specific for sample combinations with frozen gut samples (i.e. EtOH-stored feces + frozen gut, EtOH-stored gut + frozen gut) in *Gymnocypris* and *Nanorana* samples, and only two in *Triplophysa* samples (Supplementary data 4). This suggests an overall higher similarity of EtOH-stored samples, which is also evident from the NMDS graphs, especially for *Gymnocypris* and *Nanorana* (Fig. 3B). As the highest differences between sample types were observed in *Nanorana* samples, especially between frozen and EtOH-stored samples (Table 1; Fig. 3c, g), the indicator species analysis revealed a large number of OTUs that were differentially abundant in latter samples; 13 OTUs for EtOH-stored feces, 29 OTUs for EtOH-stored gut (35 OTUs for the combination of EtOH-stored samples) and 12 OTUs for the frozen gut samples (Supplementary data 4). The consistent increase of Proteobacteria in frozen *Nanorana* guts (Fig. 3A) was mainly caused by seven of these OTUs, all belonging to the Gammaproteobacteria: two undetermined Aeromonadaceae, two undetermined Enterobacteriaceae, two *Klebsiella* (Enterobacteriaceae), and *Pseudomonas* (Pseudomonadaceae). These OTUs represented very high proportion of reads in the frozen gut samples (89.7%), but most of them (9 OTUs) were completely absent from all of the EtOH-stored samples of *Nanorana*. However, the majority of these OTUs were found in low abundances in a few fecal samples of the fish hosts. Furthermore, some of the indicator OTUs for the *Nanorana* frozen gut samples were also found in high relative abundances in some of the *Gymnocypris* and *Triplophysa* frozen gut samples (Supplementary data 2, 4).

## Discussion

As the number of DNA sequences of environmental or host-associated samples in public databases increase and acquiring such data becomes a routine approach in microbial ecology, meta-analyses of comprehensive “big data” sets is becoming a promising research direction, leading to important insights into general patterns of bacterial diversity [21, 54-56]. However, considering the many technical factors influencing the outcome of amplicon analyses, including sample preservation, DNA extraction, PCR conditions and sequencing methods [57-59], it is of high importance to ascertain that such meta-analyses indeed recover biological patterns and not methodological differences among studies.

Our study exemplifies that the recovery of host-associated microbiota richness and community structure may vary among sampling substrates, but especially among commonly used sample preservation methods. The two EtOH-stored substrates, (i) gut content recovered by dissection and (ii) freshly collected feces, had differences in community structure especially in *Triplophysa* sp. (Fig. 3), but overall, revealed rather consistent biological patterns. For instance, for the two EtOH-stored substrate types, we found the highest OTU richness in *Nanorana* tadpoles, followed by *Gymnocypris*, and the lowest richness in *Triplophysa* (Figs. 1-2). This agrees with predictions derived from their diet - the largely herbivorous tadpoles have a particularly diverse microbiome [e.g. 10, 11], sub-adults of the small cyprinid *Gymnocypris* can be expected to be omnivorous [60] whereas the diet of *Triplophysa* is dominated by macroinvertebrates [61].

However, important differences between gut and fecal samples were also apparent, especially in *Nanorana* tadpoles where the relative abundance of Firmicutes was distinctly smaller in fecal samples (Fig. 3A; Supplementary data 5). This pattern was particularly driven by Clostridia of which 18 OTUs were identified as indicators, and thus were relatively more abundant in, the gut samples (Fig. 3A; Table S3). One factor affecting the community difference between gut and feces could also be the exposure to oxygen upon leaving the intestinal tract [62].

An even more divergent pattern was found for the frozen samples, with enormous differences both in bacterial richness and community structure, especially in *Nanorana* tadpoles. Consistently, in almost all individual tadpole gut samples, Proteobacteria enormously increased in relative abundance, with a large reduction of relative abundances of Firmicutes and Verrucomicrobia (Fig. 3; Supplementary data 5). As summarized by Kohl [63], blooming of certain taxa can change the composition of gut or fecal bacterial communities [40, 64], and we hypothesize this is what happened in our samples upon the short thawing periods during sample transport. It is however surprising that this blooming pattern affected the tadpole gut samples, while in the samples of the two fish, no conspicuous and consistent increase of Proteobacteria was noted, except for three *Gymnocypris* samples (Supplementary data 5). The blooming hypothesis is also supported by the fact that the Proteobacteria increase was caused by a limited number of bacterial OTUs, and most strongly influenced by only seven OTUs. Moreover, these OTUs were totally absent from the EtOH-stored *Nanorana* gut samples. Although we included a pond water control in our study to account for external contamination, these bacteria still may represent environmental contaminants from remains of pond water present on the specimens during dissection as suggested by their existence, in low relative abundances, in some of the fecal samples which were collected directly from the water.

Several studies on soil, human- and insect-associated microbiomes typically revealed differences among preservation methods being smaller than those between taxa and individuals, thus validating meta-analyses based on differently stored samples [39, 65-68]. A study about sample preservation method of fecal microbiota of spider monkeys revealed that the microbial community composition of EtOH-stored and frozen samples were similar to fresh ones [69]. Thus, it is expected that the latter sample storing methods are producing comparable results. Furthermore, a study on insect-associated microbiomes has suggested that the sample storage method (freezing, ethanol, dimethyl sulfoxide, cetrimonium bromide, storage without any preservative) has no or minor effect on microbiome composition [67]. However, under typical field sampling conditions, continuous deep-freezing of samples cannot be always ensured, which implies the possibility of radical effects on gut microbiomes. Sampling ungulate feces in the wild, Menke et al. [70] observed only moderate shifts of the microbiome during 2-4 days but radical changes afterwards, usually following rain. On the contrary, Beckers et al. [64] observed an important decrease in bacterial diversity in horse feces already after approximately 4 hours. A significant decrease of Bacteroidetes has been reported from fecal samples exposed to room temperature or to natural environmental conditions [71, 72], which we also found to be the case for the frozen samples that were exposed to thawing in this study. Our study confirms that in certain cases, a rise of “bloom” bacteria can completely obscure the original microbiome composition of fecal samples, and several bacterial families might be particularly prone to contain such rapid growth taxa; for instance, Enterobacteriaceae and Pseudomonadaceae were associated with microbiome shifts both in this study and in that of Beckers et al. [64].

## Conclusion

Our case study confirms that both substrate (gut content *vs*. feces) and preservation method can influence the analysis of intestinal microbiomes, and provides an example from aquatic vertebrates. Differences between substrates and methods are here shown for samples from exactly the same individuals, sampled at the same time point, thus excluding these factors that might influence microbiome structure. Fecal samples can be a suitable substrate to investigate intestinal microbiomes in tadpoles and fishes as their analysis recovered similar patterns of bacterial diversity as gut samples in our study. This is important because for ethical reasons, non-invasive collection of feces is to be preferred over dissection. However, when using fecal samples, one must be aware of environmental contaminations of these samples, which could be minimized by collecting control samples in the immediate environment. Yet, the substantial differences in bacterial community composition we found between gut and fecal samples indicate that these different types of substrates should only be combined with great caution in an analysis, and only when large differences between hosts are expected. The strongest differences were found between preservation methods and demonstrate that blooming of contaminant taxa can completely distort the bacterial community in samples of intestinal microbiome of aquatic vertebrates, within only a few hours of thawing as it is common under field conditions. Despite identical treatment of samples, the blooming effect in our study was much stronger and more frequent in gut samples from a largely herbivorous species (a frog tadpole), which leads to a question whether the highly diverse bacterial community of herbivores may be particularly prone to such artefacts.

## Acknowledgments

We are indebted to numerous colleagues, in particular Magnus Asmussen, Nicole Börner, Andrew Henderson, Antje Schwalb, and Anja Schwarz for their help during fieldwork; and to the team of the NAMORS station of the Institute of Tibetan Plateau Research of the Chinese Academy of Sciences, in particular Guangjian Wu, for logistic support. SA and MV were funded through the Deutsche Forschungsgemeinschaft (DFG) via the International Research Training Group 2309, “Geoecosystems in transition on the Tibetan Plateau” (TransTiP).

## Supplementary data

**Supplementary data 1.** Used primer combinations.

**Supplementary data 2.** Table of bacterial OTUs by samples.

**Supplementary data 3.** Correlation scatterplots of OTU richness (log transformed) between sample types per species (*Gymnocypris, Triplophysa* and *Nanorana*).

**Supplementary data 4.**
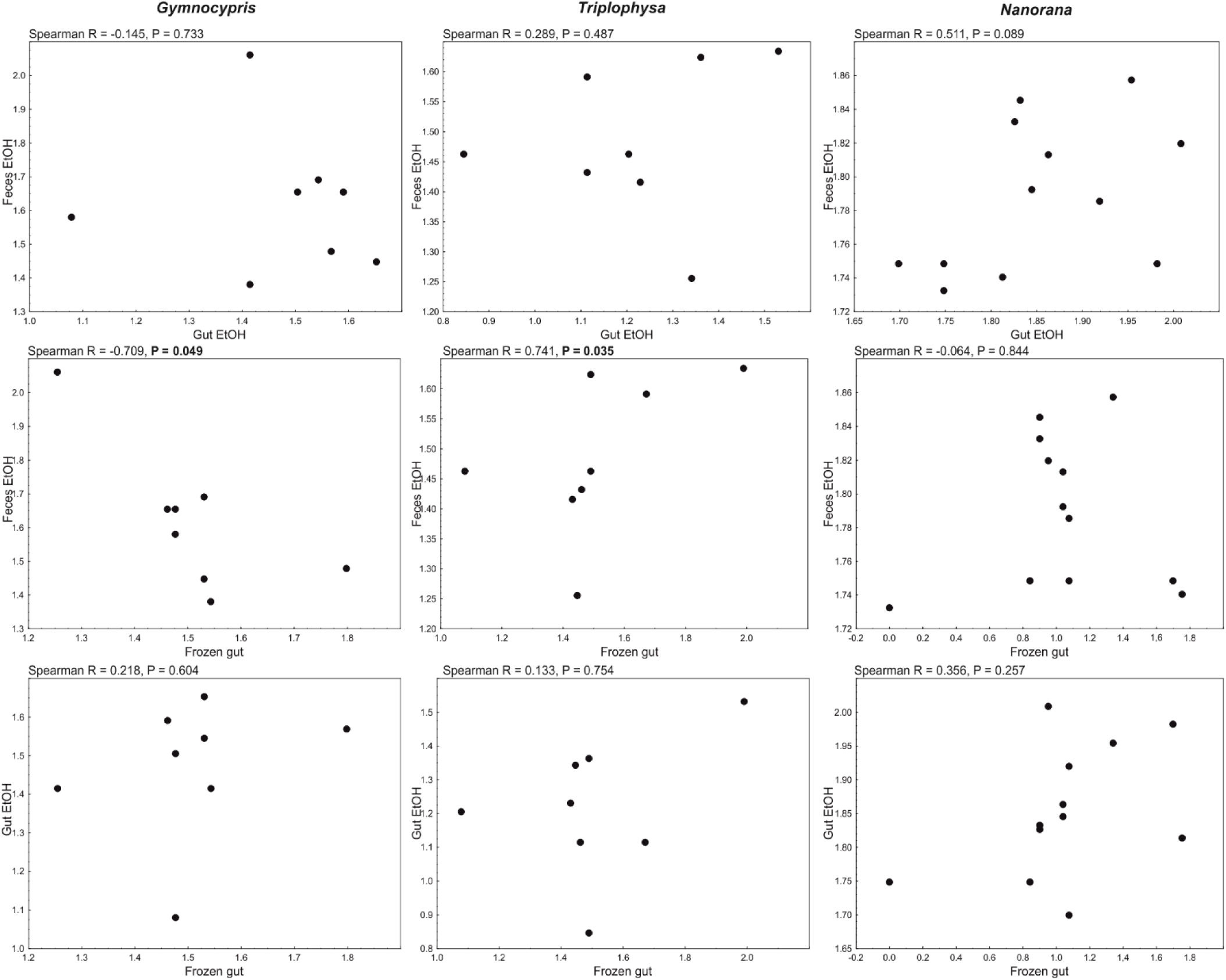
Bacterial indicator OTUs per treatment (or combination of treatments), analyzed separately per host species.

**Supplementary data 5.**
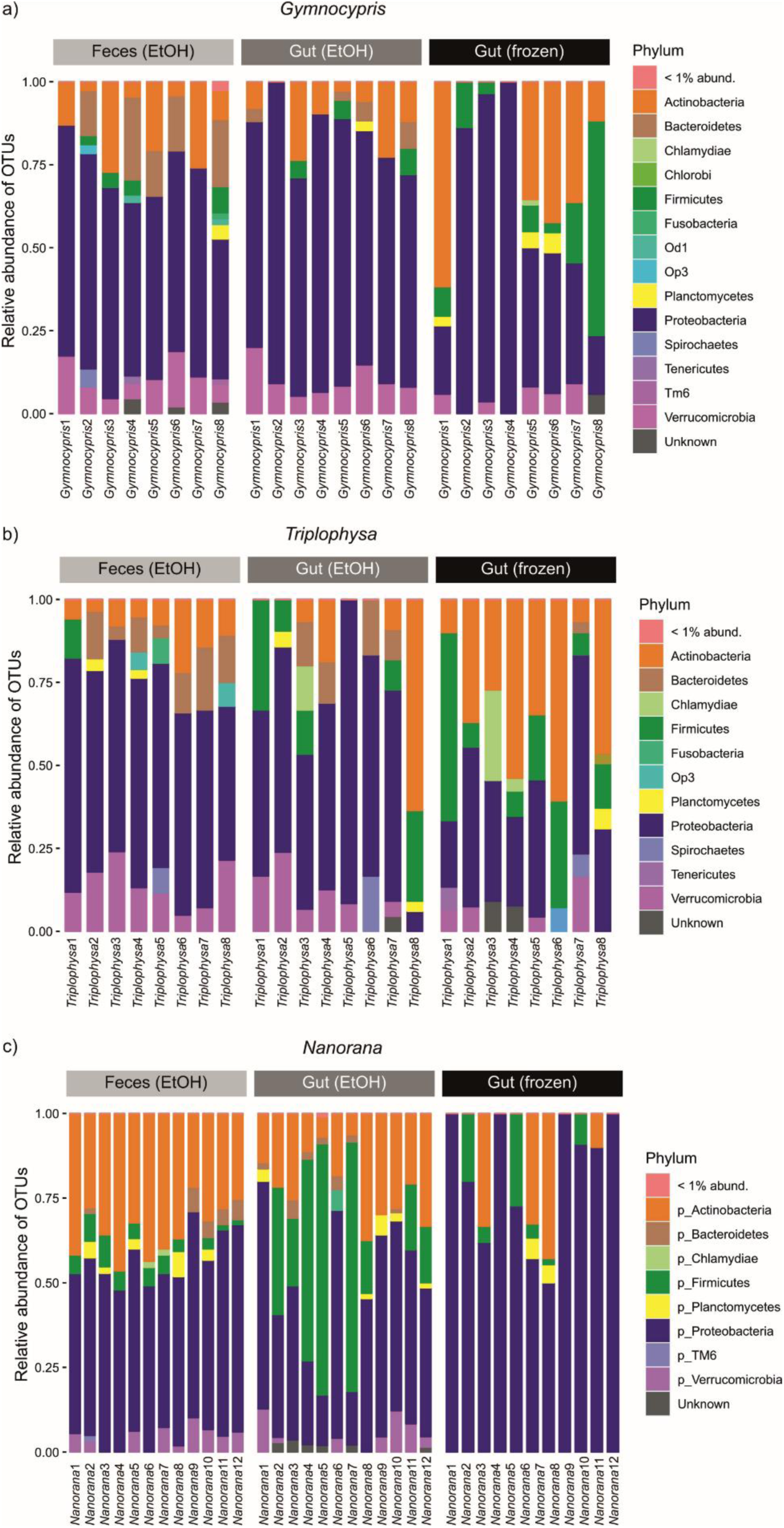
Relative abundance of OTUs in *Gymnocypris* (a), *Triplophysa* (b) and *Nanorana* (c) samples.

